# Protein-based Virtual Screening Tools applied for RNA-Ligand Docking identify new Binders of the preQ_1_-Riboswitch

**DOI:** 10.1101/2022.06.10.494309

**Authors:** Elisabeth Kallert, Tim R. Fischer, Simon Schneider, Maike Grimm, Mark Helm, Christian Kersten

**Affiliations:** Institute of Pharmaceutical and Biomedical Sciences, Johannes Gutenberg-University Mainz, Staudingerweg 5, 55128 Mainz, Germany

**Keywords:** RNA docking, RNA ligands, virtual screening, preQ_1_, Riboswitch, CADD, SBDD

## Abstract

Targeting RNA with small molecules is an emerging field. While several ligands for different RNA targets are reported, structure-based virtual screenings against RNAs are still rare. Here, we elucidated the general capabilities of protein-based docking programmes to reproduce native binding modes of small molecule RNA ligands and to discriminate known binders from decoys by the scoring function. The programmes were found to perform similar compared to the RNA-based docking tool rDOCK and the faced challenges during docking, namely protomer and tautomer selection, target dynamics and explicit solvent, do not largely differ from challenges in conventional protein-ligand docking. A prospective virtual screening with the *Bacillus subtilis* preQ_1_-riboswitch aptamer domain performed with FRED, HYBRID and FlexX, followed by microscale thermophoresis assays identified 6 active compounds out of 23 tested virtual screening hits with potencies between 29.5 nM and 11.0 μM. The hits were selected not solely based on their docking score, but for resembling key interactions of the native ligand. Therefore, this study demonstrates the general feasibility to perform structure-based virtual screenings against RNA targets, while at the same time it highlights pitfalls and their potential solutions when executing RNA-ligand docking.

## Introduction

Due to its fundamental role in many biological processes, RNA became of increasing interest as a potential drug target over the last years.^1–9^ Several antisense oligonucleotides (ASO) and small interfering RNAs (siRNA) were approved by the Food and Drug Administration (FDA) and European Medicines Agency (EMA) or are in clinical development.^10^ Targeting RNA with small molecules, however, is still a novel concept and only one molecule except of ribosome-targeting antibiotics is approved by the FDA. With Risdiplam,^11,12^ the first approved small molecule splicing modifier is available for the treatment of spinal muscular atrophy and further splicing modifiers, micro RNA (miRNA) ligands and binders of extended trinucleotide repeats are under elucidation for many indications including cancer,^8,13–16^ neurological and genetic disorders.^2,17–23^ While only around 1.5% of the human genome is transcribed into proteins, the common drug-targets, about 70-90% is transcribed into RNA.^7,24,25^ These mainly noncoding RNAs (ncRNA) represent promising targets for novel therapeutics. Further, viral RNA^26^ like the SARS-CoV-2 frameshift pseudoknot,^27,28^ enterovirus internal ribosomal entry site^29^ or the HIV *trans*-activation response element (TAR)^26,30^ were successfully targeted with small molecules. To fight emerging multidrug resistant bacteria, riboswitches (RS) are attractive targets for the development of new antibiotics.^31–33^

While structure-based computational methods like molecular docking are regularly used for protein targets, most identified RNA-binding small molecules however, emerged from high-throughput or fragment-screening campaigns, derivatisation of a known ligand or serendipity rather than structure-based design by molecular docking.^27,34–40^ One challenge in this regard is the low number of available 3D RNA structures compared to the number of proteins deposited in the protein data bank (PDB),^41^ often additionally struggling with low resolution. However, these structures already show clearly that the limited chemical diversity of just four RNA bases compared to 20 canonical amino acids in proteins can still result in many different folds, e.g. hairpins, G-quadruplexes, multi-way junctions, (pseudo)knots, L-shaped tRNAs, triple helices, ribozymes and the ribosome complex.^7^ While docking tools are occasionally used to generate an RNA-ligand binding mode hypothesis,^13,29^ prospective virtual screenings (VS)^42–44^ and extensive validation of the used docking programmes for RNA targets are rare.^45–48^ However, validation of conventional protein-based docking tools for its applicability against RNA targets is crucial because parameters cannot easily be transferred and nucleic acids hold their own unique challenges different from proteins for docking programmes, like high intrinsic flexibility, negatively charged phosphate backbone which is eventually masked by cations, or a limited chemical diversity.^49–52^ Consequently, over the time, some specialized RNA-ligand docking algorithms and scoring functions were developed.^53–62^

Once validated against a given dataset to demonstrate the ability to correctly predict crystallographic binding modes (posing) and accurately discriminate binders from non-binders or decoys by scoring, the true challenge for a docking programme is the identification of novel scaffolds for a target by VS. While for proteins prospective docking community challenges like the drug design data resource (D3R) grand challenges^63–67^ are performed from time to time, challenges for RNA structure prediction,^68–71^ but not for molecular docking have been reported so far. For our prospective case study, we selected the *Bacillus subtilis* (*Bs*) prequeosine 1 (7-aminomethyl-7-deazaguanine, preQ_1_, **1**) RS aptamer domain as a model system. RSs are noncoding, *cis*-acting RNA elements in the 5’ untranslated region (UTR) of mRNA regulating gene expression. RSs consist of a ligand binding domain and an expression platform and are usually found in prokaryotes. The preQ_1_-RS aptamer was selected due to its reported assayability,^72–74^ available crystal structures with different ligands^75,76^ and proved druggability as small drug-like ligands structurally distinct from the native ligand preQ_1_ were reported.^76^ PreQ_1_-RSs are also present in *Enterococcus faecium, Neisseria gonorrhoeae, Streptococcus pneumoniae* and *Haemophilus influenzae*, pathogens from the World Health Organization (WHO) priority list,^32^ making the preQ_1_-RS an attractive target for the development of novel antibiotics against drug-resistant bacteria.^77^

### Pose prediction

Prior prospective VS, the tools under elucidation were evaluated in terms of their ability to correctly predict native binding modes of X-ray and NMR complex structures and to discriminate known ligands from decoys with similar physicochemical properties.^78,79^ The protein-based docking programmes tested were FlexX^80,81^ and its derivative LeadIT,^82^ and FRED^83^ with its template-docking modification HYBRID.^84^ DOCK6^85,86^ was included as it was validated besides AutoDock for RNA-ligand docking previously^46,47^ and demonstrated to be successful in a prospective VS against a purine RS as well.^42,87^ rDOCK,^54^ a successor of RiboDOCK,^53^ represents a tool, developed explicitly for RNA-ligand docking. Noteworthy, all dockings were performed in a highly automated process with minimal user intervention and hence are very likely to hold optimization potential.

To elucidate the precision in pose prediction, a panel of 150 RNA-ligand complexes was compiled from previous studies^46,47,54^ including ribosome structures, riboswitches, viral RNA and artificial RNA aptamers (Table S1). Without experienced user intervention during receptor and ligand preparation prior docking, the success rates for re-docking, defined as an root-mean-square deviation (RMSD) of < 2.5 Å,^45,46,62^ was rather poor compared to what might be expected by expert-defined docking protocols. Success rates ranged from 35-50% with median RMSD values of 2.5-4.7 Å (Figure 1, Table 1). However, these success rates lie within the expected range of automated, non-optimized docking screens.^67,88^ The exception is HYBRID with a success rate of 80%, but it is to be noted that HYBRID while being based on FRED, additionally to the ChemGauss scoring function uses an additional scoring step called chemical gaussian overlay (CGO) which considers the reference ligand during docking.^84^ Therefore, a better performance for re-docking using this template-based approach is not surprising. rDOCK, which was originally developed for RNA-ligand docking, performed only slightly better than protein-based docking programmes. As recommended from the original authors, from the two scoring functions SF3 and SF5 within rDOCK, SF5 performed better, likely due to its improved desolvation terms.^54^ DOCK, which was improved and validated for RNA-ligand docking previously^47^ and was already successfully applied in a prospective VS against the adenine riboswitch,^42^ was outperformed with a success rate of only 35% and a median RMSD of 4.7 Å when using a common protein-based protocol. However, it is reported that force-field based scoring functions tend to overestimate ionic interactions.^49^ While the backbone of RNA is highly charged, these charges are neutralized or at least masked by cations in solution. For compensation, the overall backbone charges are usually modified for docking either by adding counter ions^47^ as also done in molecular dynamic (MD) simulations^89–91^ or by adaptation of the partial charges either of the backbone oxygen^42^ or phosphorous atoms.^46^ By increasing the partial charge of P by +1 to neutralize the backbone, the success rate of DOCK (“mod. charge”, Table 1) was improved by 10%.

**Figure 1.**
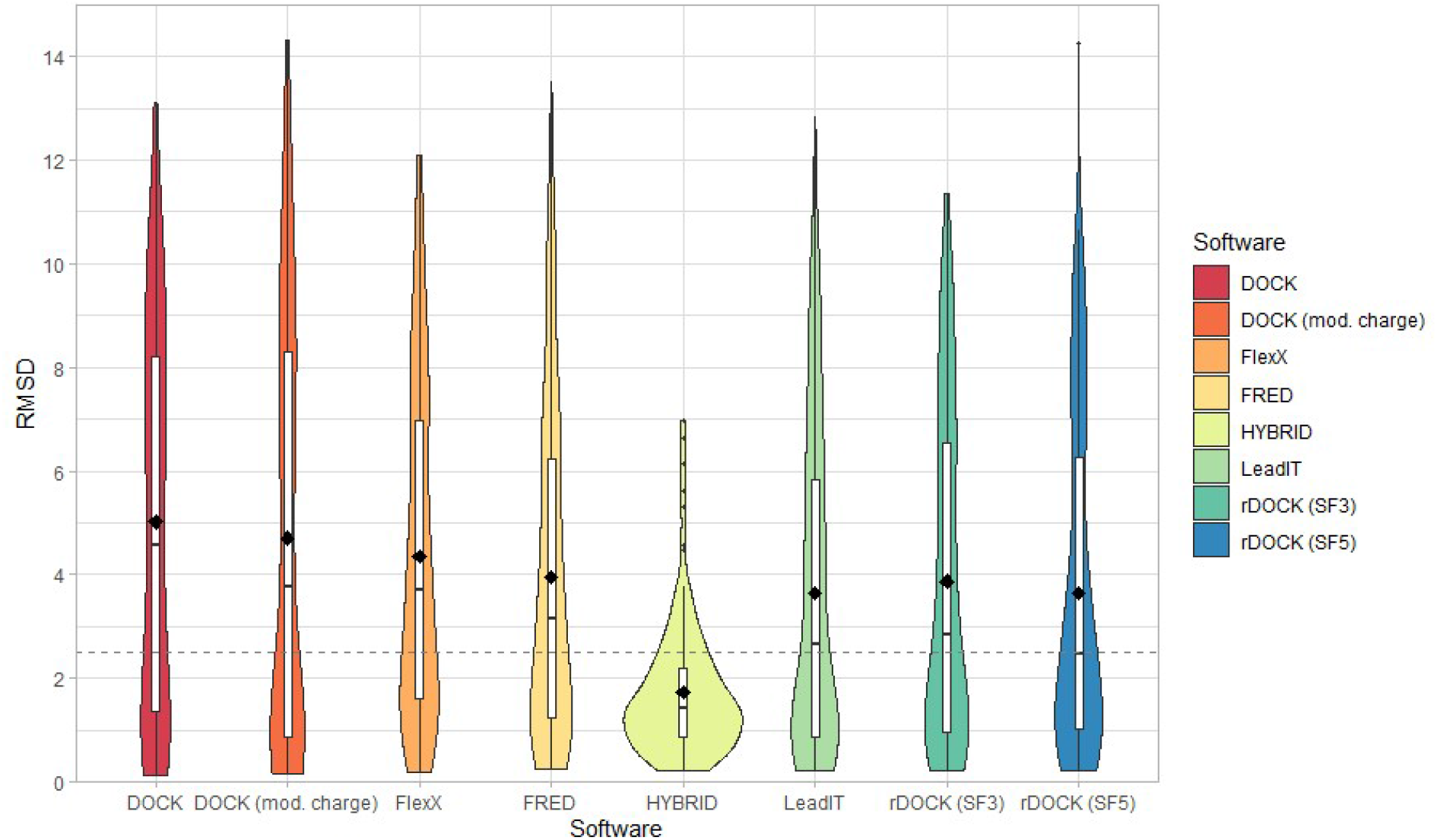
Violin plots of re-docking performance in an automated docking workflow against 150 RNA-targets for DOCK 6.9 (red), DOCK 6.9 with modified charges (orange, adding +1 partial charge on phosphorus atoms), FlexX-3 (light orange), FRED 3.3.0.3 (beige), HYBRID 3.3.0.3 (yellow), LeadIT-2.3.2 (light green), rDOCK (SF3 green, SF5 blue). Bar plots around the median are depicted with 25-75% range in white and mean value as a black dot. The dashed line at RMSD = 2.5 Å resembles the cut-off for a redocking to be defined as “successful”.

**Table 1.** Re-docking result summary of 150 RNA-ligand complexes.

| Software | Mean RMSD (Å) | Median RMSD (Å) | Success rate (%)<sup>a</sup> |
| --- | --- | --- | --- |
| DOCK 6.9 | 5.2 | 4.7 | 35 |
| DOCK 6.9 (mod. charge) | 4.7 | 3.8 | 45 |
| FlexX-3 | 4.3 | 3.7 | 40 |
| FRED 3.0.3 | 4.0 | 3.2 | 43 |
| HYBRID 3.0.3 | 1.7 | 1.4 | 80 |
| LeadIT-2.3.2 | 3.6 | 2.6 | 49 |
| rDOCK 2013.1-1 (SF3) | 3.9 | 2.9 | 46 |
| rDOCK 2013.1-1 (SF5) | 3.6 | 2.5 | 50 |

<sup>a</sup>The success rate was defined as percent of re-docking poses within an RMSD of 2.5 Å compared to the reference ligand.

### Binder-decoy discrimination

For a prospective VS, docking programmes are not only required to identify correct binding modes of known ligands, but also to be able to discriminate binders from non-binders by their docking score.^79^ While non-binders are often difficult to receive from literature, a common method is the usage of decoys that have similar physicochemical properties like the ligands, but are structurally dissimilar to be classified as non-binders. For the ribosomal 16S aminoglycoside binding site (A-site)^92^ and seven riboswitches (adenine,^42,93^ thiamine pyrophosphate – TPP,^94,95^ *S*-adenoysl methionine –SAM,^96^ flavin mononucleotide – FMN,^37,97^ preQ_1_,^73^ guanine^42^ and tetrahydrofolate – THF^98^) ligands were collected from literature and corresponding decoys were generated from the “database of useful decoys: enhanced” (DUD-E).^78^ After docking of ligands and decoys against the selected target structures, the discrimination was evaluated by receiver operating characteristic (ROC) curves and their area under the curve (AUC) for overall performance as well as the enrichment^99^ of top 2%, 5% and 10% of the screened datasets (Table 2, Fig. S2).

**Table 2.** Binder *vs* decoy discrimination performance for the ribosomal A-site and seven riboswitches.

| PDB-ID | Target | binders | decoys | ROC AUC |  |  |  |  |  |  |
| --- | --- | --- | --- | --- | --- | --- | --- | --- | --- | --- |
|  |  |  |  | (% of ligands recovered in top 2% of screened database) |  |  |  |  |  |  |
|  |  |  |  | DOCK | DOCK (mod. charge) | FlexX | FRED | HYBRI D | LeadI T | rDOC K (SF5) |
| 1LC4 | rRNA A-site | 18 | 1050 | 0.93<br>(22) | 0.91<br>(17) | 0.72<br>(6) | 0.73<br>(0) | 0.75<br>(11) | 0.68<br>(0) | 0.93<br>(44) |
| 1Y26 | adenine-RS | 13 | 900 | 0.66<sup>a</sup><br>(0) | 0.48<br>(0) | 0.97<br>(54) | 1.00<br>(100) | 1.00<br>(100) | 0.99<br>(92) | 0.99<br>(83) |
| 2GDI | TPP-RS | 9 | 550 | 0.81<sup>b</sup><br>(38) | 0.57<br>(33) | 0.67<br>(22) | 0.79<br>(33) | 0.86<br>(44) | 0.58<br>(33) | 0.86<br>(44) |
| 2GIS | SAM-RS | 3 | 150 | 0.77<br>(0) | 0.76<br>(0) | 0.88<br>(0) | 0.99<br>(100) | 1.00<br>(100) | 0.82<br>(33) | 0.92<br>(67) |
| 3F2Q | FMN-RS | 26 | 600 | 0.83<br>(12) | 0.87<br>(46) | 0.64<br>(12) | 0.88<br>(50) | 0.91<br>(58) | 0.70<br>(31) | 0.99<br>(67) |
| 3FU2 | preQ<sub>1</sub>-RS | 8 | 350 | 0.44<br>(0) | 0.44<br>(0) | 0.94<br>(75) | 0.88<br>(63) | 0.99<br>(100) | 0.97<br>(75) | 1.00<br>(87.5) |
| 4FE5 | guanine-RS | 19 | 1150 | 0.86<sup>c</sup><br>(3) | 0.58<br>(16) | 0.74<br>(8) | 0.65<br>(0) | 0.71<br>(2.5) | 0.81<br>(34) | 0.83<br>(31.5) |
| 4LVW | THF-RS | 10 | 550 | 0.40<br>(5) | 0.46<br>(5) | 0.82<br>(5) | 0.71<br>(25) | 0.73<br>(30) | 0.83<br>(10) | 0.60<br>(10) |

<sup>a</sup>Explicit water molecule ID 290 was manually included in binding site definition. <sup>b</sup>Several water molecules around reference ligand TPP were included in binding site definition. <sup>c</sup>Binding site size was reduced to 4 Å around the reference ligand.

Overall, a good discrimination between reported binders and decoys was observed with ROC AUCs over 0.6 (moderate enrichment) for all except 5 RS cases of DOCK which might require some further modification like the inclusion of explicit solvent molecules (as done for the TPP- and adenine-RS) or modification of the binding site size (as for the guanine-RS). Especially the adenine-RS is an exemplary demonstration of the importance of user intervention prior docking and screening library preparation, as the selected ligands of this RS originate from a VS with DOCK in which an explicit solvent molecule was included in the receptor preparation.^42^ For the rRNA A-site, DOCK performed better than other protein-based tools with high ROC AUCs of 0.93 and 0.91 with or without charge modification, respectively, similar to rDOCK (ROC AUC 0.93). The RNA-specialized rDOCK (SF5) showed an overall good performance for the RSs as well, with ROC AUCs between 0.60 and 1.00. However, all protein-based tools also reached high ROC AUCs and good enrichments, especially for RSs. HYBRID, due to its template-based approach with ligands often being structurally similar to the crystallographic reference ligand, demonstrated its suitability for RNA-ligand docking with ROC AUCs between 0.71 and 1.00 and recovering between 11 and 100% of the ligands within the top 2% by score of the docked datasets. The results obtained with the non-template-based tools FRED, FlexX and LeadIT were similar with only slightly reduced ROC AUCs and often reaching very good enrichments, especially for the adenine-RS, SAM-RS, preQ_1_-RS and THF-RS, all with ROC AUCs over 0.7. Hence, it can be concluded that protein-based docking tools can generally be used against RNA-targets.

### Common pitfalls and solutions

In favour of comparability between the different tools under elucidation, automated workflows for docking were established, however a more refined and user-controlled procedure of receptor and ligand preparation and docking parameters should be considered as best practice.^100–104^ While doing so for all 150 targets with all docking programmes under elucidation is beyond the scope of this manuscript, some selected cases will be discussed to highlight common pitfalls and possible solutions when performing RNA-ligand docking. While the best scoring pose did not always reproduce the native binding mode, lower scoring poses with RMSD values < 2.5 Å were regularly found for nearly all programmes and targets, indicating that correct scoring of generated poses is more challenging than coverage of generated binding complex geometries. However, in some cases docking failure can be assigned to specific features of the docking setup.

#### Tautomers

For re-docking of tetrahydrobiopterin to the THF-RS, the RMSD was 4.0 Å with a score of -28.3 kJ/mol using the automatic generated 1H tautomer with LeadIT (Figure 2 A). Manually changing to the 3H tautomer (Figure 2 B) improved the re-docking result of the top pose to an RMSD of 1.6 Å with H-bond complementarity to residues U-7, U-35 and U-42 as found in the crystal structure. The improved score of -39.4 kJ/mol further shows the ability of the scoring function to select the right tautomer over the wrong one. Similarly, the importance of tautomeric states was reported previously for the FMN-RS ligand ribocil for which the oxygen of the pyrimidinone tautomer is essential as an H-bond acceptor excluding the pyrimidinol tautomer.^34^

**Figure 2.**
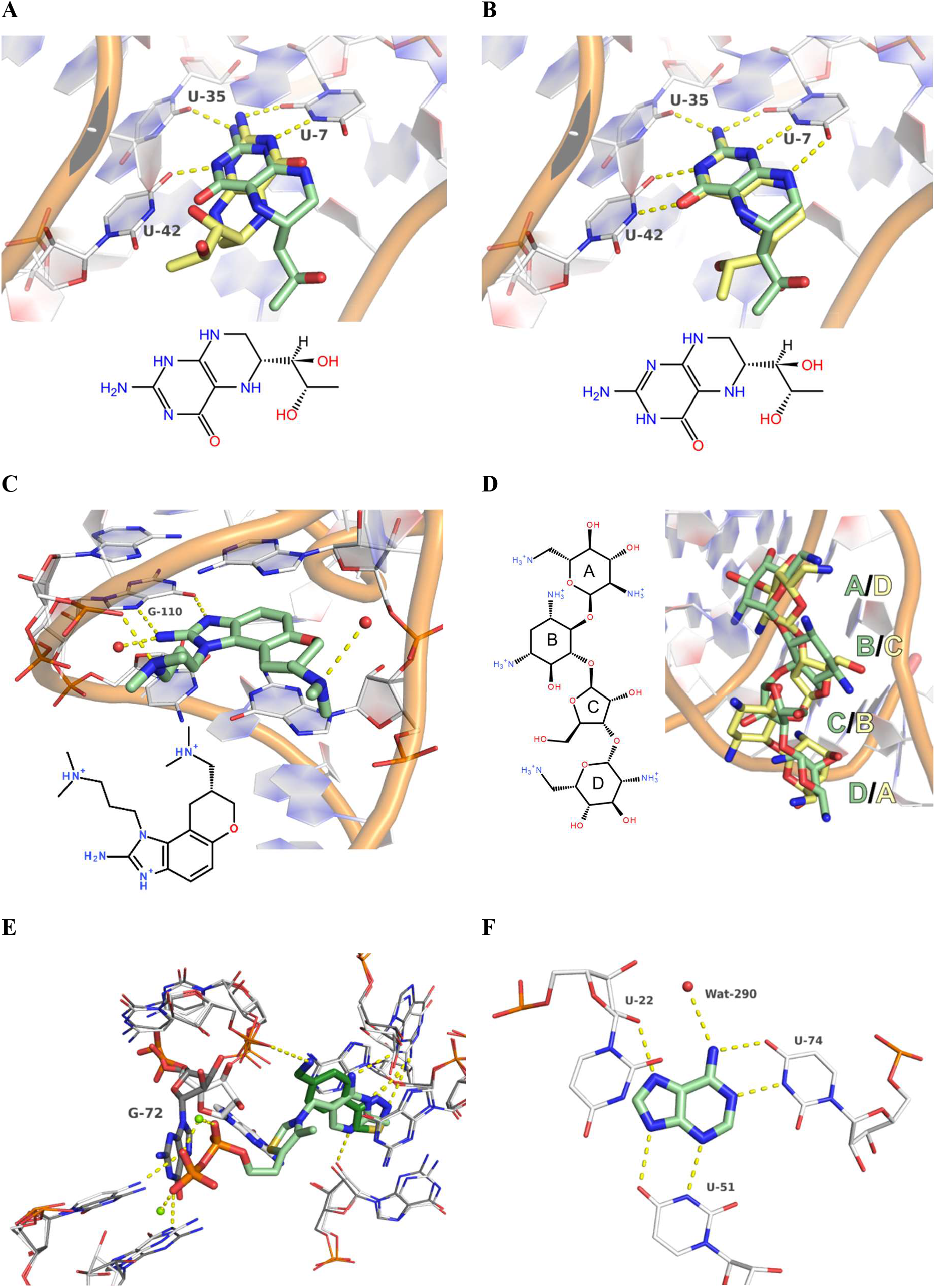
Identified challenges of RNA-ligand docking. RNA is depicted with white carbon atoms, crystallographic ligands with pale green carbon atoms, docking poses with light yellow carbon atoms and polar interactions as yellow dashed lines. **A**) Predicted binding mode of the tetrahydobiopterin 1H-tautomer in the THF-RS (re-docking RMSD 4.0 Å, FlexX-score -28.3 kJ/mol, PDB-ID 4LVW). **B**) Predicted binding mode of the tetrahydobiopterin 3H-tautomer in the THF-RS (re-docking RMSD 1.6 Å, FlexX-score -39.4 kJ/mol, PDB-ID 4LVW). **C**) Crystallographic binding mode of 2-aminobenzimidazole in its three-fold protonated state forming two H-bonds as a donor to G-110 (PDB-ID 3TZR). **D**) Predicted binding mode of neomycin in complex with HIV TAR (PDB-ID 1QD3) reveals an inverted binding mode of the docking pose compared to the NMR structure, however displaying highly similar interactions. **E**) Superposition of the TPP-RS in complex with TPP (white RNA carbon atoms and pale green ligand carbon atoms, PDB-ID 2GDI) and a fragment ligand (grey RNA carbon atoms and dark green ligand carbon atoms, PDB-ID 4NYB). G-72 adopts an altered conformation depending on which ligand is bound. **F**) Adenine binding to its RS (PDB-ID 1Y26). Inclusion of Wat-290 is crucial to improve binder-decoy discrimination with DOCK.

### Protomers

For re-docking of a synthetic ligand to the hepatitis C virus internal ribosomal entry site (HCV IRES, PDB-ID: 3TZR),^105^ re-docking with FRED resulted in an RMSD of 5.5 Å (score -11.2 kcal/mol). In this case the ligand was protonated by Protoss^106^ prior docking, resulting in a two-fold protonated molecule with an uncharged 2-aminobenzimidazole moiety. However, two crucial H-bonds with G-110 of HCV IRES require the protonated state of this substructure (Figure 2 C). With a predicted pK_a_ of 8.4 (MarvinSketch-18.13.0, ChemAxon Ltd., http://www.chemaxon.com), the relative occurrence of the +3 protomer is about 90% at physiological pH. Docking of this protomer resulted in a correct pose with RMSD = 1.5 Å and slightly better score (−12.0 kcal/mol). Interestingly, docking of the deprotonated ligand resulted in RMSD values between 0.92 and 2.14 Å for rDOCK, LeadIT and FlexX (Table S1), indicating that the wrong protonation state can be compensated by other interactions and only FRED being sensitive to the correct protonation state. Also, different protonation states of multi-basic ligands like aminoglycosides should be considered. While aliphatic amines are clearly basic and protonated under physiologic conditions if observed alone, the relative occurrence of protomers might be of importance in RNA-ligand recognition.^107^ Exemplarily, the six-fold protonated state of neomycin is dominant at pH 7.4, but with only 22% (calculated with MOE, Figure 2 D). Several minor species with lower formal charges are also possible under these conditions and potential binding protomers of the target RNA or changes of protonation states might occur upon binding.

### Beyond “rule of five” ligands

Aminoglycosides hold further challenges by being large and polar, exceeding the (extended) Lipinski’s rule of five (RO5) for drug-like, orally bioavailable molecules.^108^ Keeping in mind that molecular docking is often applied in VS to identify lead structures, molecular docking programmes are often trained with drug-like molecules. As within the re-docking set of 150 RNA-ligand complexes only 46 ligands do not violate the RO5, the programmes are not only challenged by the target RNA, but also by many of the ligands. Especially for aminoglycosides like neomycin with a molecular weight of 621 g/mol, 25 H-bond donor and 19 H-bond acceptor functionalities, a tPSA of 363 Å^2^ and 16 rotatable bonds, docking predictions are challenging. Considering the linear construction of four ring systems however, even high RMSD re-docking poses might be considered as no absolute failure. The roughly symmetric neomycin is simply inverted within the binding site, but aliphatic rings are fairly aligned and polar interactions reasonably saturated as observed for the LeadIT-predicted pose in complex with the human immunodeficiency virus *trans*-activation response element (HIV TAR) (Figure 2 D, PDB-ID 1QD3).^109^ While the RMSD for the re-docking pose is 9.5 Å, pseudo similarity (a metric within LeadIT to describe nearest atoms of same type instead of the identical atom ID) is 1.9 Å.

### Target dynamics

Many RNA molecules are of highly dynamic nature.^49–51^ Hence, docking against a single conformation might not be sufficient to represent RNA-ligand binding behaviour. While re-docking was always preformed against frame 1 of the NMR structures, this might not necessarily be the best frame selection method even though frame number 1 usually represents the lowest energy conformation.^46^ Other strategies might be the selection of the frame closest to average coordinates or docking against an ensemble of frames either derived from NMR structures or MD simulations. When docking against all frames of 11 NMR complex structures for which multiple frames were deposited in the PDB, the mean re-docking RMSD was reduced from 6.6 Å to 3.7 Å (Table S2). Noteworthy, for some complexes the ligand RMSD compared to average coordinates reached values up to 5.2 Å over the different NMR frames, highlighting the high flexibility even within the bound state of the complex. Additionally, it is to mention that docking against NMR structures is usually referred to be less precise and accurate compared to docking against X-ray structures.^46^ Besides intrinsic dynamics, distinct conformational changes of residues, as reported for the TPP-RS, might need consideration. In the complex of several fragments with the TPP-RS, G-72 adopts a different orientation compared to the TPP- and analogue-bound state (Figure 2 E).^95,110^ To consider this conformational change, target flexibility should be taken into account during docking either by a protocol allowing side-chain flexibility^62^ or a strategy with an ensemble of multiple RNA-structures.^111,112^

#### Solvation

Lastly, solvation and desolvation effects upon ligand binding and water-mediated interactions should be elucidated. While explicit solvation sites within protein binding pockets can have an huge impact on affinity and selectivity,^113,114^ it is more than likely that similar effects account for RNA binding sites as well. However, within the RNA-ligand docking test set presented, only 18 structures with a resolution below 2.0 Å allow sufficiently accurate determination of explicit water molecules. For the adenine riboswitch a structurally important water molecule (PDB-ID 1Y26)^115^ was reported previously (Figure 2 F).^42^ Including this water molecule in the docking setup of the binder-decoy discrimination of DOCK improved the ROC AUC from 0.47 to 0.66 (Table 2). Similarly, the enrichment for the TPP-RS was also improved by inclusion of crystallographic solvent molecules.

#### Further considerations

While we identified the importance of the correct selection of ligand protomers and tautomers (Figure 2 A-C), similarly rare protomers and tautomers of RNA might also play a role for ligand recognition in specific cases.^116,117^ Additionally, it is to be taken into account that besides the four nucleobases, over 170 RNA modifications are reported^118–120^ which occur in biological systems and require individual tautomer and protomer considerations and potentially parameterization.^121^

### Virtual screening against the preQ_1_-riboswitch

Due to their overall good performance, the fast protein-based docking programmes FlexX, FRED and HYBRID were selected for a prospective VS against the *Bacillus subtilis* preQ_1_-RS. All three programmes showed high accuracy in binding mode prediction with re-docking RMSD values of 0.8, 0.4, 0.5 Å and binder-decoy discrimination with ROC AUCs of 0.94, 0.88 and 0.99, respectively (Table S1, Table 2). Further, not only the discrimination of binders from non-binders, but also the ranking within a series of 9 reported ligands^73^ spanning four orders of magnitude in potency, showed reasonable correlation between docking scores and reported pK_D_-values (Figure 3 A, Table S3). Likewise, a correlation between the number of matched interactions resembling those of the native ligand preQ_1_ with its RS (Figure 3 B) and the pK_D_ was observed for the 9 molecules as well. These interactions include three Watson-Crick-like H-bonds with C-15, additional H-bonds with U-6 and A-30 and possible H-bonds between the basic amine and G-5 and the phosphate backbone. Further, preQ_1_ stacks between G-11 and G-5 with face-to-face π-π-interactions. From this interaction profile, a pharmacophore model containing these seven features nicely correlating with affinity (Figure 3 A/B) was derived.^75^ For VS, our in-house VS-library of 3.34 million compounds was filtered for physicochemical criteria (see material and methods) and the remaining 12,507 compounds were docked without any constraints. Molecules for purchase were selected based on their docking scores and matching of the pharmacophoric features for at least one of the three programs, including molecules with (1) high scores and many met pharmacophore features, (2) high scores and little reflection of native ligands’ interactions valuing the docking score over “MedChem intuition”, as well as (3) low scoring molecules that resemble many pharmacophore features valuing human decision making over the scoring function. Subsequently, 23 compounds were purchased for testing (Table S4). Due to previous filtering, the selected molecules were not flagged as a *pan*-assay interference compounds (PAINS).^122^ However, as a note of caution PAINS definition for RNA binding, might differ from conventional protein-PAINS definition, e. g. multi-basic *pan*-RNA binders spermine and spermidine^123^ are also not flagged as PAINS and the concept of promiscuous binding might need further elucidation in the context of RNA ligands.

**Figure 3.**
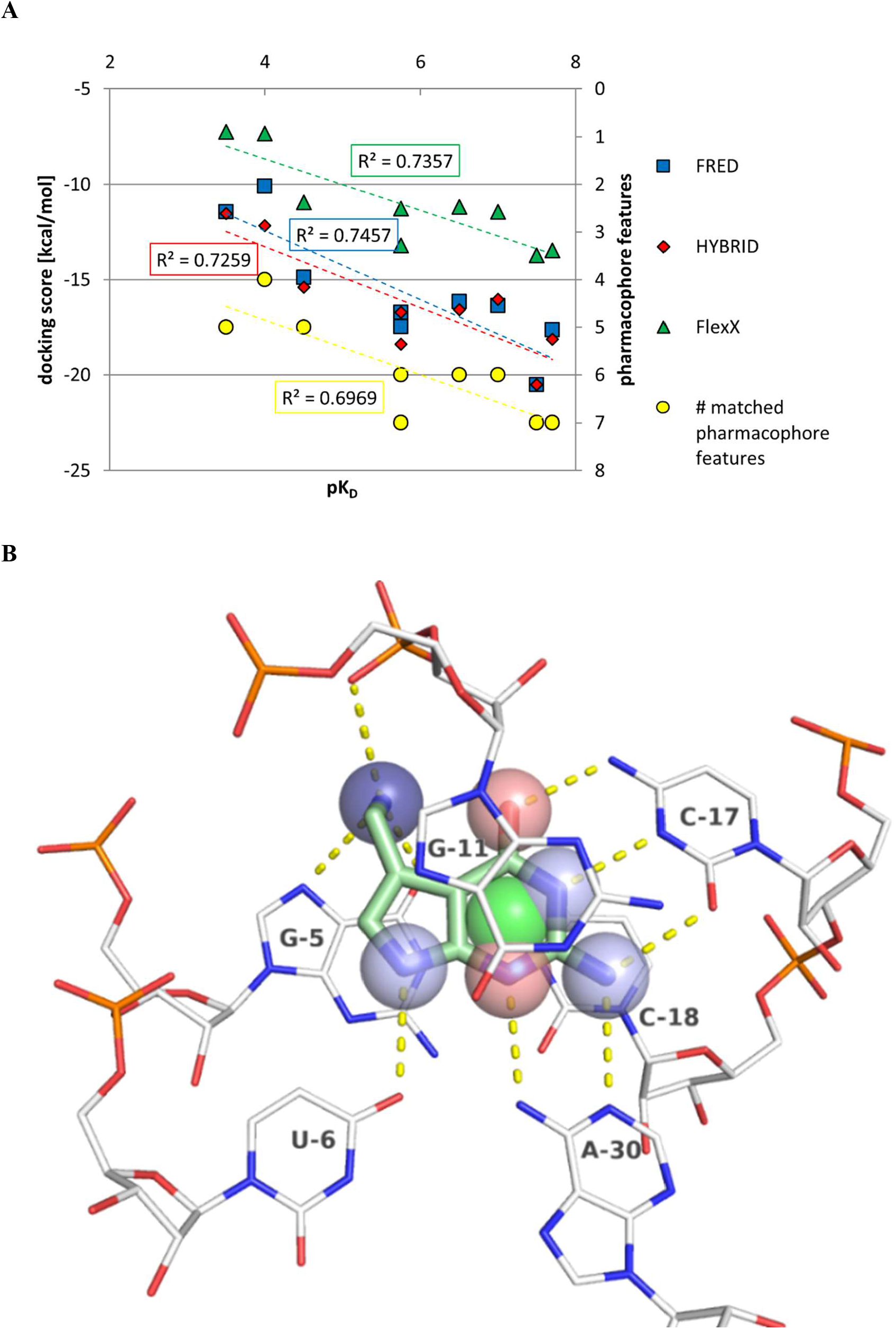
**A**) Correlation plots between docking score (left y-axis) and reported pK_D_ values for protein-based docking tools FRED (blue), HYBRID (red) and FlexX (green) against the preQ_1_-RS and between number of matched pharmacophoric features (right y-axis, yellow) and pK_D_. **B**) PreQ_1_ (pale green carbon atoms) in complex with the preQ1-RS (white carbon atoms) including polar interactions (yellow dashed lines) and pharmacophore features described in the text: dark blue = H-bond donor or cation, light blue = H-bond donor, salmon red = H-bond acceptor, green = aromatic center, PDB-ID 3FU2.

### Experimental results

PreQ_1_-RS aptamer binding of the purchased compounds was elucidated by microscale thermophoresis (MST) using a Cy5-labelled RNA. The assay was validated by K_D_-value determination of the reported^72,73^ preQ_1_-RS ligands preQ_1_, preQ_0_, guanine, 7-deazaguanine and 7-methylguanine (**1-5**, Figure 4 A) with K_D_ values similar to reported results (Table S3). For VS hits, a binding pre-screening using three different concentrations of 100 μM, 31.6 μM and 10 μM was performed. For 6 out of 23 tested compounds, **6**-**11**, which showed a concentration-dependent shift in thermophoresis, K_D_ values were subsequently determined (Figure 4 B, Figure S3). Compounds **6, 7, 8, 10** and **11** matched 5-6 pharmacophore features of preQ_1_ (Figure 3 B) and additionally showed high docking scores in at least one of the three docking setups (FlexX, FRED, HYBRID, Table S4), while **9** matched 6 of 7 pharmacophore features with a comparatively low score. The conserved interactions according to the generated docking poses among all newly identified binders include the Watson-Crick like H-bonds with C-15, H-bonds with A-30 and aromatic stacking with G-11 (Figure 5 A-D). Lastly, none of the molecules with high docking scores and a low number of matched pharmacophore features turned out to bind to the preQ_1_-RS (Table S4). Hence, the importance of proper pose inspection after docking is highlighted^124^ even though the structural diversity and novelty is thus limited. This especially accounts for **6** (the strongest binder 8-oxoguanine) and **7** (8-aminoguanine) being highly similar to previously reported ligands. The exceptional high potency of **6** with a K_D_ of 29.5 nM compared to **2**-**4** and **7** however, cannot be easily assigned to additional direct H-bonds to the RNA based on the predicted binding mode (Figure ***5*** A). Potentially water-mediated interactions which require an H-bond acceptor at the 8-position are formed. This could also explain the much lower potency of the close homolog **7** with a K_D_ of 9.07 μM which carries an H-bond donor at the corresponding position but obtains very similar docking scores. Structurally most distinct from preQ_1_ and other guanine-derivatives, **8** still matches H-bond patterns with C-15 and A-30 but lacks the H-bond donor to U-6 according to docking poses predicted by FlexX and HYBRID. This compound however is positively charged which might compensate for the missing H-bond with U-6 and it might also form additional polar or hydrophobic/aromatic contacts (Figure 5 B). **9** is the smallest binder identified in this VS and showed the lowest docking scores being ranked 1326^th^, 5948^th^ and 398^th^ out of 12,507 for FlexX, FRED and HYBRID, respectively. However, based on the predicted binding modes, this small molecule still matches 6 out of 7 pharmacophore features of preQ_1_ (Figure 5 C) in all docking setups, only lacking one H-bond donor/cationic moiety, resulting in a K_D_ of 2.01 μM. Lastly, for tetra- and 7,8-dihydrobiopterin (**10** and **11**) binding with K_D_ values of 2.26 and 11.0 μM, respectively, a crucial role of the H-bond donor to interact with U-6 (Figure 5 D) was indicated. For the aromatic pterins biopterin and neopterin (**12** and **13**, Table S4) which lack the hydrogen bond donor functionality at the corresponding position, no binding was detected in the MST assay even though predicted binding modes had high scores and still matched 5 of the pharmacophore features. Similarly, this was observed for 7-methylguanine (**5**) previously, which binds less strongly to the preQ_1_-RS compared to **3** and **4** (Figure 4 A).^73^

**Figure 4.**
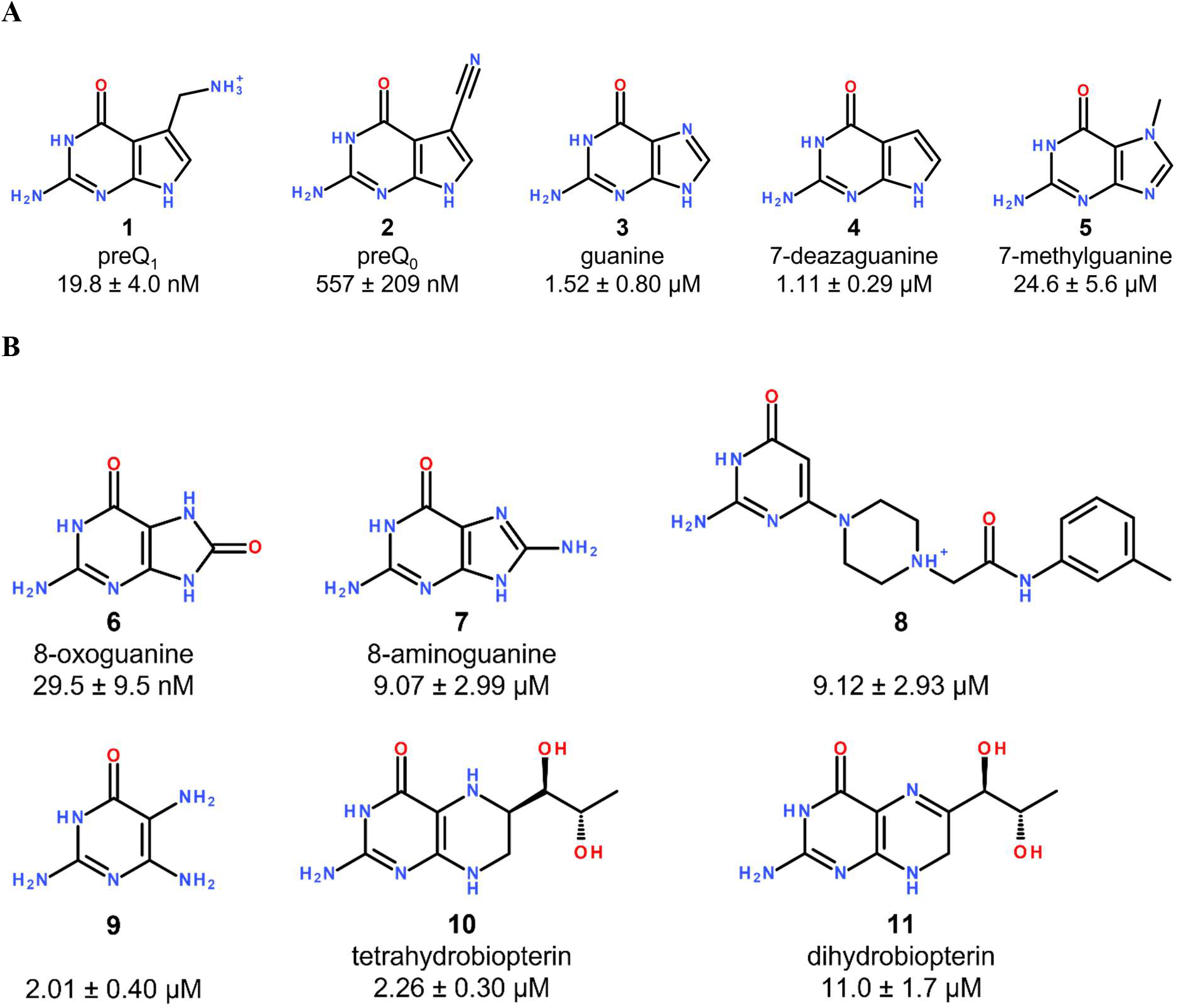
MST-derived K_D_ values for known preQ_1_-RS ligands (**A**) and new ligands identified *via* VS (**B**).

**Figure 5.**
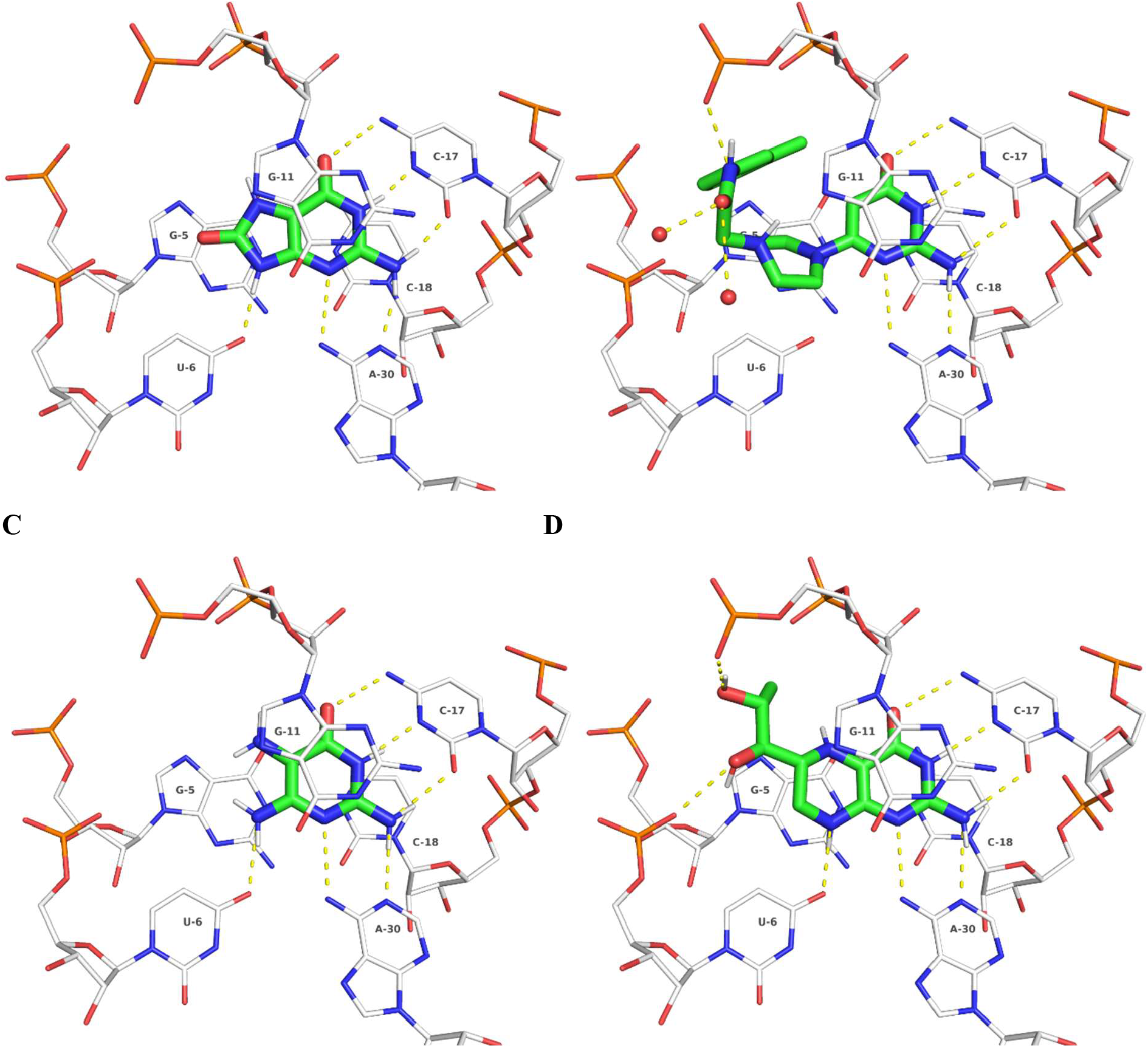
Predicted binding modes of **6** (**A**), **8** (**B**), **9** (**C**) and **10** (**D**) in complex with the preQ_1_-RS aptamer (PDB-ID 3FU2). For clear view only the predicted binding modes of the FlexX-docking are shown. Ligands are depicted with green carbon atoms including polar hydrogens, RNA with white carbon atoms. H-bonds are shown as yellow dashed lines. H-bonds known from preQ_1_ (Figure 3 B) with C-15 and A-30 as well as aromatic stacking with G-11 are conserved among all ligands and one H-bond with U-6 for all except **8**.

## Conclusion

Targeting RNAs with small molecules is an emerging field, but structure-based approaches are limited to very few cases due to a lower number of known RNA-ligand complexes compared to proteins. Hence, the knowledge basis for molecular docking is also restricted and RNA holds its own challenges for docking-based VS. The basic physical principles of molecular recognition however, do not differ between RNA and proteins and molecular docking is applicable. Using a compilation of 150 RNA-ligand complexes (Figure 1, Table 1), the capability to reproduce native binding modes for protein-based docking tools without major modifications was similar to or even better than DOCK which was improved^47^ and successfully used^42^ for RNA-ligand docking previously and competitive to the RNA-specific rDOCK.^54^ Additionally, the ability to discriminate known ligands from decoys for the rRNA A-site and several riboswitches was demonstrated (Table 2). The faced challenges for RNA-ligand docking include the selection of correct protomers and tautomers, target flexibility and (de-)solvation effects, all of which are known for proteins as well, and are not RNA-specific (Figure 2). Subsequently, in a prospective VS with the preQ_1_-RS aptamer, 6 new ligands with K_D_-values between 29.5 nM (**6**) and 11.0 μM (**11**) were discovered (Figure 4 B). Based on the hit-selection criteria it can be concluded that a good docking score only in combination with binding mode interpretation based on a pharmacophore hypothesis was superior to the docking score alone as no molecule with a high score and low number of matched pharmacophore features was identified as a preQ_1_-RS binder (Table S4). In lieu thereof, ligand **9** (K_D_ = 2.01 μM) matched 6 of 7 pharmacophore features with a relatively low score. Consequently, we conclude that RNA-ligand docking with conventional software is generally feasible but requires a certain extend of human intervention in both docking setup validation and hit selection. Hence, the used strategy herein may be used as a guideline to identify and circumvent potential pitfalls and support structure-based VS against RNA targets in the future.

## Material and methods

### Molecular docking and structure-based virtual screening

RNA-ligand complex structures were obtained from the PDB.^41^ For pose-prediction, PDB entries from previous publications^46,47,54^ were combined and visually inspected using PyMOL.^125^ If multiple chains of the target structure were present in the asymmetric unit of X-ray structures, chain A was selected. In case of NMR structures, frame 1 usually representing the lowest energy conformation^46^ was used. If ions were present within the binding site, these ions were kept during docking. Except for LeadIT, which by default includes water molecules forming at least three interactions with receptor and ligand, water molecules were removed from the complexes. All docking programmes under elucidation treated the ligand to be flexible and the receptor to be rigid. If not stated differently, only the best scoring pose was further evaluated.

For LeadIT-2.3.2 docking,^82^ the graphical user interface (GUI) was used for receptor preparation using standard settings and a binding site definition including residues within 6.5 Å around the reference ligand, which was protonated within LeadIT using the Protoss module.^106^ The docking was performed under default settings using the enthalpy/entropy hybrid approach with 200 iterations per placement and fragmentation.

For FlexX^80,81^ docking, the receptor and ligand were exported from LeadIT and docking was performed without any further modifications from command-line.

For FRED/HYBRID docking^83,84,126,127^ the receptor was prepared using the Make_Receptor functionality of the OpenEye tools under default conditions. Ligand conformers for docking were generated using OMEGA^128,129^ based on the Merck Molecular Force Filed (MMFF94).^130^ For macrocyclic ligands, OMEGA with the keyword “macrocycle” was used. Docking with FRED and HYBRID was subsequently performed without deviations from default settings.

Receptor preparation for rDOCK-2013.1-1^54^ was performed using the molecular operating environment (MOE 2015).^131^ For rDOCK, a 6.0 Å binding site was defined around the reference ligand and 50 poses were generated and subsequently scored using the SF3 and SF5 scoring functions.

For DOCK 6.9^85,86^ a protocol based on the tutorial of the developers was followed with only minor modifications. The receptor and ligand were protonated and AM1-BCC ligand charges were applied within MOE 2015. To define the ligand binding site, a grid exceeding the ligand by 6.0 Å with a grid spacing of 0.3 Å was generated and receptor parameters were obtained from the AMBER99 force field. The extra margin of the receptor box was 5.0 Å. A second docking protocol was performed adding an additional charge to the phosphate backbone of the RNA to obtain a formal charge of zero per nucleotide to mimic counter ions masking the formal charge (phosphorous atom partial charge was changed from 1.1662 to 2.1662 as described previously).^46^ Due to a poor performance under these settings in some cases of the binder-decoy discrimination set, explicit solvent molecules were included for docking against the adenine- and TPP-RS and the binding site was reduced to 4.0 Å around the reference ligand for the guanine-RS.

For the prospective VS against the preQ_1_-RS, PDB-ID 3FU2,^75^ chain A was used. Docking with FlexX-4.1,^132^ FRED/HYBRID-3.3.0.3 was performed as described above. Our in-house virtual molecule library of trusted vendors derived from ZINC15^133,134^ contained 3.34 million molecules with at most 1 violation of Lipinski’s RO5,^108^ no reactive and no PAINS^122^ moieties, a tPSA below 200 Å^2^ and less than 11 rotatable bonds. This library was filtered after protonation with MOE 2019 using OpenEye FILTER to include only molecules without known aggregator functionalities, 4-9 H-bond donors, 2-10 H-bond acceptors, a molecular weight between 130 and 400 g/mol, xlogP between -3 and +3, tPSA over 50 Å^2^, 0-8 rotatable bonds, a formal charge of 0-2 and less than 4 chiral centres to reduce chemical complexity. The remaining 12,507 molecules were subsequently energetically minimized using OMEGA-classic for FlexX docking or conformers were generated by OMEGA-pose for FRED/HYBRID docking. After docking without constraints, poses were sorted by docking score and elucidated by visual inspection and their accordance to the preQ_1_ pharmacophore features (Figure 3 B) within MOE 2019 prior selection for purchase (Table S4). Some structural analogues of promising scaffolds, including **7** (analogue of **6**), **10** and **11** (analogues of **12, 13** and **29**, Table S4) not being part of the initial screening library, were docked and added for testing separately after the initial VS.

### Microscale thermophoresis

The HPLC-purified, Cy5-labeled, *Bs*preQ_1_-RS aptamer, sequence 5’-Cy5-AGAGGUUCUAGCUACACCCUCUAUAAAAAACUAA-3’ was purchased from Eurofins genomics, Germany GmbH. MST experiments were performed on a Monolith Pico (NanoTemper Technologies, Munich, Germany) using standard uncoated capillaries. The RNA was diluted to 20 nM in 50 mM Tris-HCl buffer (pH 7.5) containing 100 mM KCl, and 25 mM MgCl_2_ similar to a protocol described previously.^72^ The preQ_1_-RS was heated to 75 °C for 5 min and cooled down to room temperature over 60 min. Ligands were added to final concentrations of 100, 31.6 and 10 μM in the initial screening. If a concentration-dependent shift of thermophoresis was observed, for K_D_-determination concentrations ranging from 1.0 mM (if permitted by solubility, otherwise starting from 316 or 100 μM) to 0.1 nM in a 3.16-fold (half-logarithmic) dilution series and one DMSO negative control without ligand were prepared. Final DMSO concentration was always 2%. All measurements were performed at least as triplicates. Results were analysed from signals after 1.5 s laser on-time with the MO.Affinity Analysis software, version 2.3 using the K_D_-fit model. Identities and purities > 90% for all compounds (Table S4) were stated in the vendor’s certificates of analysis (CoA). For all newly discovered preQ_1_-RS binders **6-11**, identities and purities >97% were confirmed by HPLC-ESI/MS (Figure S4). The mass spectra were obtained from a 1100 series HPLC system from Agilent with a Poroshell 120 EC-C18 150 × 2.10 mm, 4 μm column. The mobile phase consisted of 80% aceonitrile, 10% H_2_O and 10% of a 0.1% solution of formic acid in water. The UV detection wavelength was 254 nm. The molecular mass was detected using an Agilent 1100 series mass-selective detector (MSD) trap with positive mode electron spray ionization (ESI). The LC−MS chromatograms and their corresponding mass spectra were analyzed using MestReNova (v.12.0.4) from Mestrelab Research.

## Supporting information

Compound SMILES

binders and decoys (sdf)

docking poses (sdf)

## Acknowledgement

We thank Prof. Tanja Schirmeister for scientific discussion and proof-reading of the manuscript and Jessica Emsermann for support during elaboration of the MST assay. The MST instrument was provided by the Bundesministerium für Bildung und Forschung (BMBF) to M.H. (BMBF / 01ED1804). The project was funded by the Internal University Research Funding (Stufe-I) of the Johannes Gutenberg-University. We further thank OpenEye Scientific and the Kuntz Lab (UCSF) for free academic licenses.

## Abbreviations

A-site: aminoglycoside binding site
ASO: antisense oligonucleotide
AUC: area under the curve
*Bs*: *Bacillus subtilis*
CGO: chemical gaussian overlay
DUDE: database of useful decoys enhanced
EMA: European Medicines Agency
ESI: electron spray ionization
FDA: Food and Drug Administration
FMN: flavin mononucleotide
GUI: graphical user interface
HCV: hepatitis C virus
IRES: internal ribosomal entry site
MD: molecular dynamic
miRNA: micro RNA
MSD: mass-selective detector
ncRNA: non-coding RNA
PAINS: *pan*-assay interference compounds
PDB: protein data bank
preQ_1/0_: prequeosine-1/0
RMSD: root-mean-square deviation
RO5: rule of five
ROC: receiver operating characteristic
RNA: ribosomal RNA
RS: riboswitch
SAM: S-adenoysl methionine
siRNA: small interfering RNA
TAR: trans-activation response element
THF: tetrahydrofolate
TPP: thiamine pyrophosphate
UTR: untranslated region
VS: virtual screening
WHO: World Health Organization

## Data and Software Availability

Molecular dockings were performed with LeadIT, FlexX, DOCK, FRED, HYBRID and rDOCK using the cited versions. Additionally, FILTER, Make_receptor and OMEGA from the OE-toolkit and MOE were used for the described steps in receptor and ligand preparation. Used PDB entries for re-docking, ligands and decoys (derived from the DUD-E webservice) for discrimination, SMILES of newly discovered preQ_1_-RS ligands with their predicted binding modes are provided in the Supporting Information.

## TOC graphic

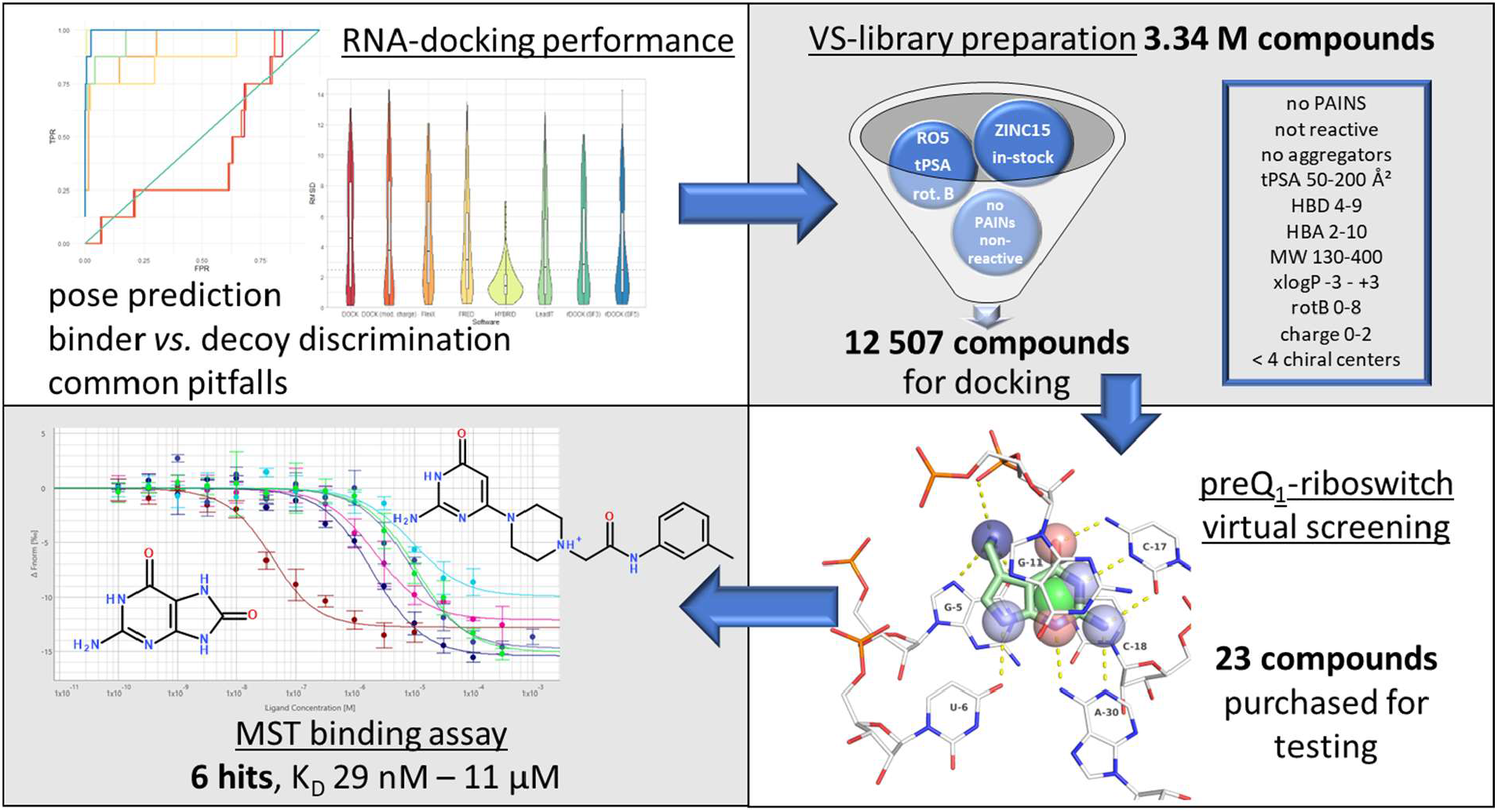

